# fMRI detects bilateral brain network activation following unilateral chemogenetic activation of direct striatal projection neurons

**DOI:** 10.1101/741710

**Authors:** Yuki Nakamura, Yukari Nakamura, Assunta Pelosi, Boucif Djemai, Clément Debacker, Denis Hervé, Jean-Antoine Girault, Tomokazu Tsurugizawa

## Abstract

Although abnormal structural and functional connectivity in the striatum during neurological disorders has been reported using functional magnetic resonance imaging (fMRI), the effects of cell-type specific neuronal stimulation on fMRI and related behavioral alterations are not well understood. In this study, we successfully combined DREADD-technology with fMRI (chemo-fMRI) to investigate the alterations of spontaneous neuronal activity induced by the unilateral activation of dopamine D1 receptor-expressing neurons (D1-neurons) in the mouse dorsal striatum (DS). We compared the effects of two different DREADD ligands, clozapine (CLZ) and clozapine-N-oxide (CNO), on behavior and fMRI. We found that the effects of CLZ on behavior were more rapid than those of CNO. In fMRI, both systemic CLZ and CNO administrations, which evoked unilateral activations of D1-neurons in DS, increased the fractional amplitude of low frequency fluctuations (fALFF) in the thalamus and bilateral cortex. In addition, we found the increased gamma-band of local field potentials in DS and bilateral cortex after CLZ-evoked unilateral activation of D1-neuron in the striatum. These results provide bases for better interpretation of cell type-specific activity changes in fMRI.

## Introduction

Pathological changes in the striatum contribute to a wide variety of neurological disorders including Parkinson’s disease (PD) (Albin et al., 1989). The dorsal striatum (DS) is the main input nucleus in the basal ganglia circuit, receiving afferents from cerebral cortex and thalamic nuclei, and plays a central role in motor control (Tepper et al., J, 2007, Belin et al., 2009). The medium-size spiny neurons of the DS are inhibitory GABAergic spiny projection neurons (SPNs), correspond to ~95% of the striatal neurons and form the sole striatal output. The rodent DS SPNs are schematically categorized into two populations according to their projection sites and the genes they express: the dopamine D1 receptor-expressing striatonigral SPNs (D1-SPNs) and D2 receptor-expressing striatopallidal SPNs (D2-SPNs) (Gerfen et al., 2000, Valjent et al., 2009). D1-SPNs project directly to the substantia nigra and the internal globus pallidus (GPi), forming the direct pathway, and their inhibitory effect on tonically active nigro-thalamic and pallido-thalamic GABA neurons leads to disinhibition of the thalamus. The activation of the direct pathway is thought to promote motor behavior (Nelson and Kreitzer, 2014). Although abnormal structural and functional connectivity in the striatum in the context of neurological disorders including in PD patients has been reported using magnetic resonance imaging (MRI) (Rauch et al., 1997), the consequences of activation of selective neuronal subtype on functional MRI (fMRI) are not well understood.

Recently, new experimental approaches for manipulating specific neurons have been developed, including optogenetics with channelrhodopsins and chemogenetics with designer receptor exclusively activated by designer drugs (DREADDs) (Roth BL et al., 2016). The DREADDs, which are genetically-modified G-protein-coupled receptors, induce relatively long-lasting (min-hours) neuronal activation, which can induce dynamic network changes. The activation of D1-SPNs in the DS by optogenetics was reported to activate a wide range of brain regions (Lee et al., 2016, Bernal-Casas et al, 2017). Chemo-fMRI with D1-specific neuronal activation has not yet been investigated.

In the present study, using fMRI in mice, we studied brain-wide functional activity resulting from D1-SPN-specific activation in the DS by human M3 muscarinic DREADD (hM3D). DREADDs were selected to be unresponsive to endogenous ligands and activated selectively by pharmacologically inert clozapine-N-oxide (CNO) (Roth BL et al., Neuron, 2016). However, it was recently shown that clozapine (CLZ) back-metabolized from CNO in vivo, can be a better stimulant than CNO itself in vivo. CLZ binds to DREADDs with higher affinity than CNO and shows a higher permeability across the blood brain barrier (Gomez et al., 2017). In this study, we aimed (1) to investigate the alteration in behavior and spontaneous neuronal activity by unilateral activation of DS D1-SPNs using the chemo-fMRI system and (2) to compare the effects of CNO and CLZ. Since striatal dysfunctions in fMRI were reported in several human neurological pathologies, such as PD or Huntington’s disease, and since D1- and D2-specific abnormalities are implicated in mouse models of these diseases (Fieblinger et al., 2014, Ferguson et al., 2011, Alcacer et al., 2017), a better understanding of relationships between cell type-specific activity and fMRI will provide insight into the analysis of these disorders, potentially leading to the development of better diagnosis and treatment for human diseases.

Fractional ALFF (fALFF) of the fluctuation of the resting state fMRI signals, which measures the relative contribution of low frequency band (0.01 – 0.1 Hz) to whole frequency range, is shown to reflect spontaneous neural activity of the brain at basal state. The functional connectivity is studies to investigate the synchronization of the BOLD fluctuation between spatially separated regions, while fALFF allows to investigate the amplitude of regional neuronal activity. In the present study, we used fALFF to map the altered neuronal activity in the whole brain. Then electrophysiology was used to optimize the altered power of the neuronal activity in the regions where fALFF was changed following the DREADD-induced neuronal activity.

## Method

### Experimental subjects

In this study, we used male and female heterozygous BAC Drd1∷Cre (D1-Cre) transgenic mice (GENSAT project; EY262 line expressing Cre recombinase under the control of the dopamine D1 receptor gene (*Drd1*) promoter). Mice were on a C57BL/6J genetic background and approximately 12-week-old at the beginning of experiments. We also used 12-week-old C57BL/6JRj mice purchased from Janvier Laboratories (France). Mice were housed under a 12-hour light/12-hour dark cycle with free access to food and water. The mice were injected adeno-associated virus (AAV) vectors in left DS and were subjected to behavioral tests and fMRI. After the experiments, mice were sacrificed and used for immunohistofluorescence (**Fig. 1A**). For fMRI study, sixteen mice were allocated to two groups: DREADD group (DREADD, n=8) and C57BL/6 group (C57BL/6J, n=8). The DREADD group was comprised of D1-Cre mice unilaterally injected with Cre-dependent DREADD-AAV and the C57BL/6 group was comprised of control C57BL/6JRj mice which received CNO or CLZ to evaluate the DREADD-independent effects of these drugs. Each group of mice was treated with CNO (DREADD-CNO or C57BL/6-CNO), CLZ (DREADD - CLZ or C57BL/6-CLZ) or saline (DREADD-saline or C57BL/6-saline) and the rsfMRI images were acquired after administration at indicated time on a different day. All animal procedures used in the present study were approved by the *Ethical committee for animal experiments* (*Comité d’Ethique en Expérimentation Animale*, *Commissariat à l’Energie Atomique et aux Énergies Alternatives, Direction des Sciences du Vivant*, Fontenay-aux-Roses, France) and by the *Ministère de l’Education Nationale de l’Enseignement Supérieur de la Recherche* (France, project APAFIS#8196-201704201224850 v1). All methods in this study were performed in accordance with the relevant guidelines and regulations.

**Figure1.**
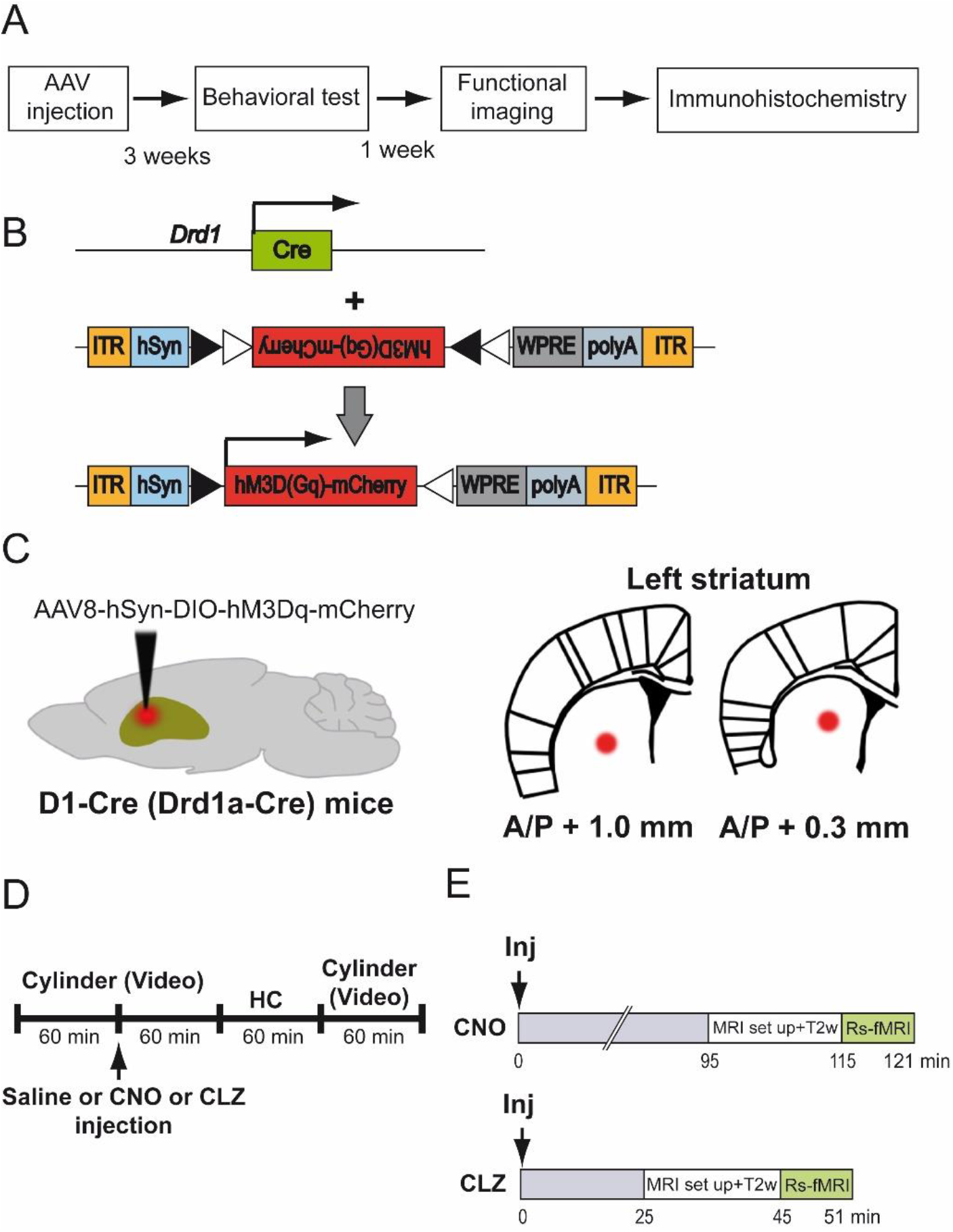
Experimental design and DREADD expression. **A.** Schematic experimental design. **B.** Schematic representation of Drd1a^Cre^ allele and hM3D (Gq)-mCherry-expressing AAV construct employing the DiO (**d**ouble floxed **i**nverse orientation) systems [pAAV8-hSyn-DIO-hM3D(Gq)-mCherry]. ITR; inverted terminal repeats. hSyn; human synapsin promoter. WPRE; woodchuck post-transcriptional regulatory element. **C.** Scheme illustrating AAV injection site in the striatum. Cre-inducible AAV vector coding for the hM3D (Gq) was unilaterally injected into the left DS of D1-Cre transgenic mice. A/P; anterior/posterior distance from bregma (mm). **D.** Schedule of behavioral test. Mice were administered vehicle, CNO (2 mg/kg) or CLZ (0.1 mg/kg) 60 min after introduction in the test chamber (cylinder). Mice were returned to their home cages 60 min after injection and were placed back in their respective cylinder 120 min after injection. Mice were video-recorded in the cylinder and locomotion and number of rotations were evaluated. HC; home cage **E.** fMRI experimental paradigm. Vehicle, CLZ (0.1 mg/kg) or CNO (2 mg/kg) were administered to the mice in the home cage. Mice were then anesthetized with isoflurane just before MRI scanning and 4-min T2 weighted image followed by 6-min resting state fMRI was scanned 95 min after CNO injection and 25 min after CLZ or saline injection. T2w; T2-weighted image.

### Viral vector injections

Mice were anesthetized with a mix of ketamine (80 mg/kg) and xylazine (10 mg/kg) and mounted on a stereotaxic apparatus (Kopf, France). Cre-inducible AAV vectors coding for the Gq-coupled DREADD, AAV8-hSyn-DIO-hM3D (Gq)-mCherry (titer ≥ 4×10^12^ vg/mL; University of North Carolina Vector Core, Chapel Hill, NC and Addgene, Cambridge, MA), were unilaterally injected into the left DS of D1-Cre mice (**Fig. 1B** **&** **C**). Two injections (0.5 μl each) in the left DS were performed at the following coordinates based on the Allen Brain Atlas: AP 1.0 mm, ML −1.5 mm, DV −3.3 mm and AP 0.3 mm, ML −2.0 mm, DV −3.5 mm from the bregma (**Fig. 1C**). All injections were performed using a 10 μl Hamilton Neuros 33G syringe at a flow rate of 0.2 μl/min. The needle was left in place for 2 min before and 5 min after the injection and then slowly retracted. Mice were placed in their home cage for more than 2 weeks for recovery.

### Rotation test

The number of 360° rotations and horizontal locomotion were measured three weeks after surgery. To assess turning movements and locomotion, mice (n=8) were placed individually inside a transparent plastic cylinder (20 cm diameter, 15 cm height) 60 min before the drug or saline administration and video-recorded. They were then injected with saline (0.9% NaCl solution, i.p.), CNO (2 mg/kg, i.p., ENZO Life sciences) or CLZ (0.1 mg/kg, i.p., Sigma-Aldrich) and video-recorded for 60 additional minutes. The dose of CNO and CLZ was determined from previous reports (Gomez et al., 2017, Alexander et al., 2009). Mice were returned to their home cages for 60 min and then (i.e. 120 min after injection) they were placed back in their respective cylinder, where they were video-recorded for another 60 min (**Fig. 1D**). Video recordings were used to visually count the number of clockwise (right, contralateral rotations to the virus-injected side) and anticlockwise (left, ipsilateral rotation to the virus-injected side) full turns. The mice were tested for 4 consecutive days: the first 2 days, vehicle or CLZ were injected in a counter-balanced design, 24 h apart and the next 2 days, vehicle or CNO were injected in a counter-balanced design, 24 h apart. Data were collected in 10-min bins.

### fMRI

fMRI acquisition was conducted in a Bruker 7T system with a mouse brain array coil (Bruker BioSpin, Ettlingen, Germany). Each mouse was lightly anesthetized with isoflurane (1.5% for induction and 0.8-1.0% for maintenance) in air containing 30% O_2_. The head was fixed with ear-bars and a teeth-bar. The fMRI acquisition started around 10 min after switching to 0.8-1.0% isoflurane. The respiratory rate was between 80-120/min as measured with a small pneumatic pillow placed next to the animal abdomen (model 1025; SA Instruments, NY, USA) and rectal temperature was maintained at 37°C during the measurement by circulating hot water. Scanning was started 25 min following saline (0.9% NaCl solution) or CLZ (0.1 mg/kg in saline) i.p. injection, and 95 min following CNO injection (2 mg/kg, i.p., **Fig. 1E**). Each mouse received CNO, CLZ or saline on a different day and the injection sequence was randomized. The dosage of CNO and CLZ was the same as for behavioral tests. The scanning time was determined from the results of rotation behavior in the cylinder test. After correction of the field homogeneity, structural T2-weighted images were acquired for 4 min. Then the rsfMRI acquisition was launched. The rsfMRI images were acquired using a T2*-weighted multi-slice gradient-echo EPI sequence with the following parameters: TR/TE = 2,000/12 ms, spatial resolution = 150 × 150 × 500 μm^3^, 180 volumes (total 6 min). rsfMRI acquisition was started at 45 min and 115 min following CLZ and CNO injection, respectively. For normalization in image processing, the structural image with the same field of view as the fMRI was acquired using T2-weighted multi-slice rapid acquisition with relaxation enhancement (RARE) with the following parameters: TR = 2,500 ms, effective TE = 13 ms, spatial resolution = 100 × 100 × 500 μm^3^, RARE factor = 4, and 4 averages.

### Image processing

The rsfMRI data analysis toolkit (REST1.8, Lab of Cognitive Neuroscience and Learning, Beijing Normal University, China) and SPM8 software were used to analyze the fMRI data and to perform the preprocessing, including the slice timing correction, motion correction by realignment, co-registration and normalization. Before preprocessing, template image co-registered to the Allen mouse brain atlas was obtained (http://atlas.brain-map.org/). The frame-wise displacement (FD) was used to check the head motion (Power et al, NeuroImage, 2012). The FD was calculated by the 6 motion parameters (3 translation and 3 rotation) from realignment in SPM. All subjects were confirmed that the head motion was less than the criteria: (1) mean FD averaged in all time-point during the scanning is less than 0.02 mm and (2) FDs in all time-point are less than 0.05 mm. There was no significant difference of mean FD among all groups. The functional and structural images were normalized to these template images. The preprocessed fMRI data was then detrended and slow periodic fluctuations were extracted using a bandpass filter (0.01 – 0.1 Hz). Mean signals in the ventricles and the white matter, and six motion parameters of an object (the translational and rotational motions) were regressed-out from the time-series of each voxel to reduce the contribution of physiological noise, like respiration and head movement. The residuals resulting from this regress-out were then smoothed with a Gaussian filter (0.3 × 0.3 mm^2^ in each slice). The smoothed images were used for all subsequent analyses in fractional amplitude of low frequency fluctuation (fALFF).

### fALFF

The fALFF was defined as the ration of total power within the frequency range between 0.01 and 0.1 Hz and total frequency range in each voxel to generate an fALFF map for each mouse. The fALFF maps were computed in each animal and then significant differences of fALFF among the groups were tested voxel-by-voxel using one-way analysis of variance (ANOVA) test with standard progressive matrices with a threshold of p < 0.05 (FDR at cluster level). Since there were no significant differences in ALFF between the 4 groups (DREADD-saline, C57BL/6-CNO, C57BL/6-CLZ, and C57BL/6-saline groups, data not shown), we used these 4 groups as a control group compared to the DREADD-CNO or DREADD-CLZ group.

### Electrophysiology

Electrophysiological recordings were performed separately outside of the MRI bore (Tsurugizawa et al., 2019). The animals, first anesthetized with 1.5% isoflurane, were placed in a stereotaxic frame (David Kopf instrument, CA). Body temperature was maintained at 37°C using a heating pad (DC temperature controller; FHC Inc., Bowdoin, ME, USA). The skull was exposed and a hole (1 mm diameter) was drilled to insert the micro-electrode. The tungsten microelectrodes (< 1.0 MΩ, with 1 μm tip and 0.127-mm shaft diameter, Alpha Omega Engineering, Nazareth, Israel) was positioned in the left DS (AP 0 mm, ML −1.8 mm, DV −3.5 mm from the bregma) or the left/right motor cortex (AP 0 mm, ML ±1.5 mm, DV −1.5 mm from the bregma). After surgery, isoflurane concentration was changed to 0.8 - 1.0%, which was the same concentration as for the fMRI experiment. The electrode was connected to a differential AC amplifier Model 1700 (AM systems, Sequim, WA, USA), via a Model 1700 head stage (AM systems, Sequim, WA, USA). Local field potentials (LFPs) were continuously recorded at 10 kHz sampling rate using dedicated data acquisition software (Power Lab, AD Instruments, Dunedin, New Zealand). CLZ or saline was injected 5 min after the start of the recording. Recording was then continued for 51 min (total recording time, 56 min). Total LFP recording was constructed with two sessions. Saline was injected in the first session and CLZ was injected in the second session. The reference electrode was positioned on the scalp.

### Band limited power (BLP) analysis

From LFP signal, five BLP time series were calculated: delta (1-4 Hz), theta (4-8 Hz), alpha (8-12 Hz), beta (18–30 Hz) and gamma (60–100 Hz) using PowerLab (AD Instruments, Dunedin, New Zealand) (Thompson et al., 2013, Tsurugizawa et al., 2019). The mean power of each frequency band in the left (ipsilateral to the DREADD-expressing side) DS and in the left and right motor cortex was calculated during 45 - 51 min after the start of injection of saline or CLZ.

### Immunofluorescence

Mice were rapidly deeply anesthetized with pentobarbital (500 mg/kg, i.p., Sanofi-Aventis, France) and perfused transcardially with 4% (weight/vol) paraformaldehyde in 0.1 M sodium phosphate buffer 90 min following CNO administration. Brains were post-fixed overnight at 4°C and cut into free-floating sections (30 μm) with a vibrating microtome (Leica) and kept at −20°C in a solution containing 30% ethylene glycol (vol/vol), 30% glycerol (vol/vol) and 0.1 M phosphate buffer. Immunolabeling procedures were as previously described (Valjent et al., 2000). After three 10-min washes in Tris-buffered saline (TBS, 0.10 M Tris and 0.14 M NaCl, pH 7.4), free floating brain sections were incubated for 5 min in TBS containing 3% H_2_O_2_ and 10% methanol, and rinsed again in TBS 3 times for 10 min. Brain sections were then incubated for 15 min in 0.2% (vol/vol) Triton X-100 in TBS, rinsed 3 times in TBS and were blocked in 3% (weight/vol) bovine serum albumin in TBS, and then incubated overnight at 4°C with primary antibody diluted in the same blocking solution. Sections were then washed three times in TBS for 15 min and incubated 1 h with secondary antibodies. After washing again, sections were mounted in Vectashield (Vector laboratories). Primary antibodies were rabbit antibodies against c-Fos (1:1000; Santa Cruz Biotechnology, #sc-52), chicken antibodies against mCherry (1:1500; Abcam, #ab205402). Secondary antibodies were anti-rabbit A633 antibody (1:400, Invitrogen, #A-21070) and anti-chicken Cy3 antibody (1:800; Jackson Immuno Research, #703-165-155). Images were acquired using confocal microscope with a 20X objective.

## Results

### Validation of DREADD expression in the striatum

We confirmed the successful AAV-transduction and hM3D-mCherry fusion protein expression using immunostaining (**Fig. 2**). Immunostaining for mCherry showed that the DREADD-expressing area was located in the injected left DS. No signal was detected in the right DS. We tested the functionality of the receptor by immunostaining for c-Fos, a marker of activated neurons, 90 min following CNO administration. A pronounced cFos immunoreactivity was observed in the left DS as compared to the right DS. The DREADD expression areas of individual mice used for fMRI are shown in **Supplementary Fig.1**. Medial part of DS was overlapped in all mice (n=8).

**Figure 2.**
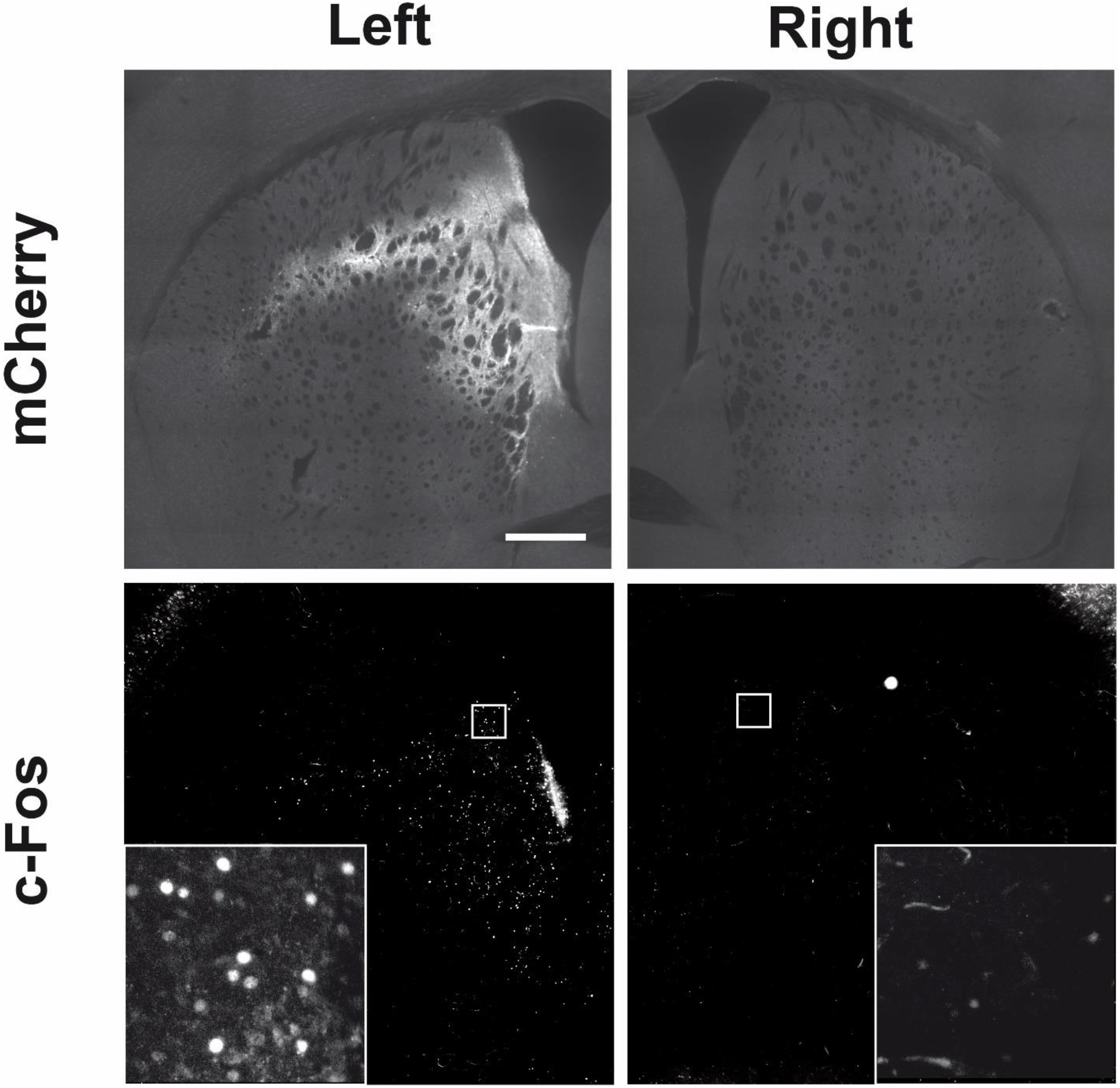
Representative images of DREADD expression. mCherry and c-Fos immunostaining in the striatum of a representative mouse. Cre-inducible AAV vectors coding for the Gq-coupled hM3D [pAAV8-hSyn-DIO-hM3D(Gq)-mCherry] was unilaterally injected into the left DS of D1-Cre transgenic mice. Higher-magnification images are shown in c-Fos immunostaining images. Scale bar, 500 μm.

### Behavioral effects of DREADD stimulation of left D1-SPNs

We examined the effects of chemogenetic D1-SPNs activation on locomotion and whole-body movements by counting the number of left and right rotations. D1-Cre mice that had been injected with AAV8-hM3D (Gq)-mCherry in the left DS 3 weeks before, were tested in a cylinder following CNO (2 mg/kg), CLZ (0.1 mg/kg) or saline administration. CNO significantly increased the locomotion 0-60 min and 120-180 min after injection, while CLZ significantly increased the locomotion 0-60 min, but not 120-180 min, after injection compared to saline injection (**Fig. 3A, 3B**). Both CNO and CLZ significantly increased the rotational bias toward the right side (contralateral to the DREADD-expressing side) compared to saline injection. The time point of maximum behavioral change was 130-140 min following CNO (2 mg/kg) injection and 20-30 min following CLZ (0.1 mg/kg) injection **(Fig. 3C, 3D)**. These data showed that CLZ (0.1 mg/kg) increased the number of contralateral rotations much more and earlier than CNO (2 mg/kg).

**Figure 3.**
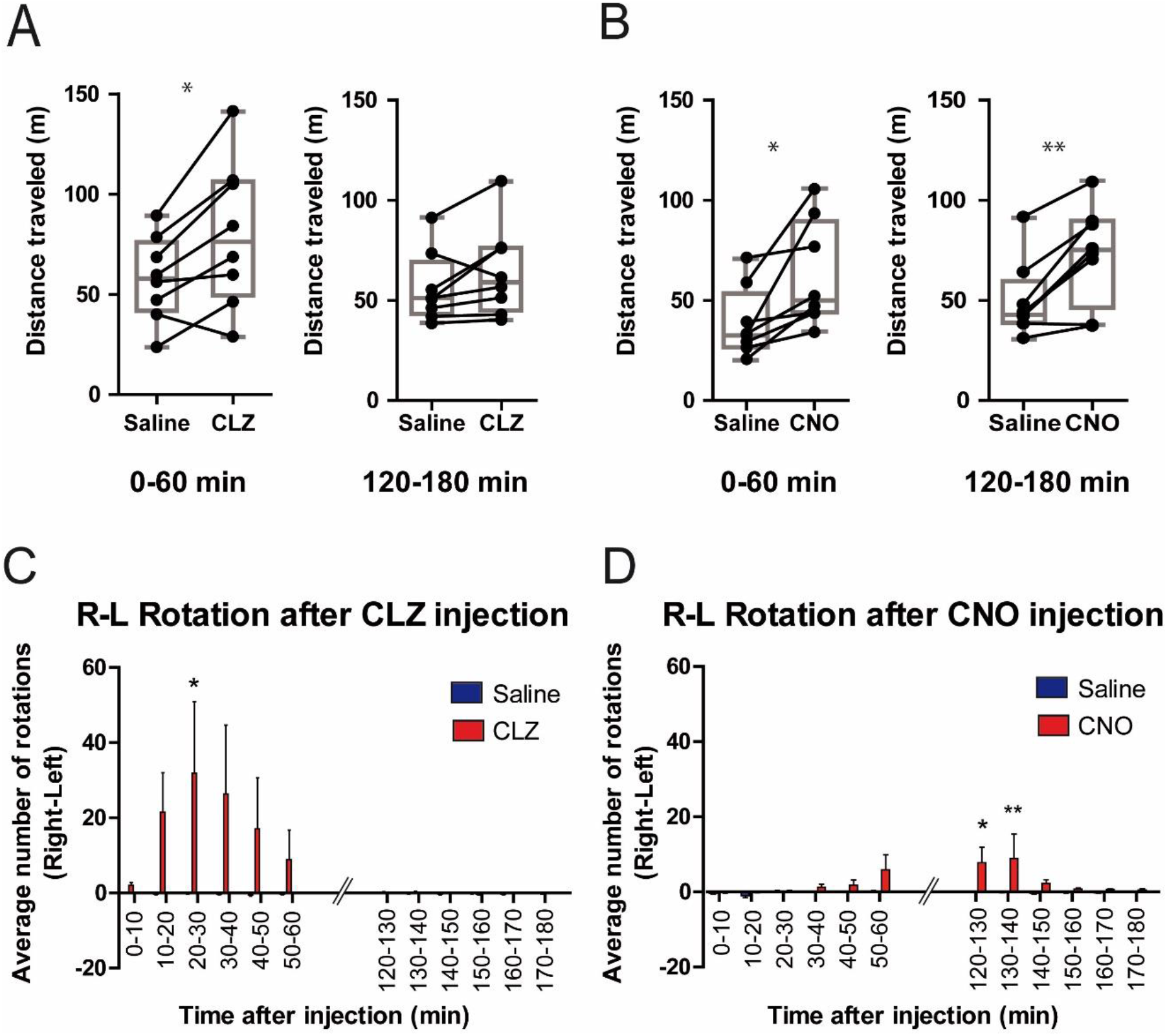
Unilateral Gq-DREADD activation in D1-SPNs induces contralateral rotations. Mice were unilaterally injected in the left DS with AAV expressing Gq-DREADD only in D1-SPNs. Mice were placed in a cylinder, injected with CNO, clozapine (CLZ) or saline and their locomotion and rotations were measured during the periods indicated in Fig. 1D. **A. B.** Locomotor activity was measured 0–60 min and 120-180 min after CLZ (**A**) or CNO (**B**) injection compared to saline administration. **C.** CLZ (0.1 mg/kg) induced contralateral rotation asymmetry [Number of full turns per 10 min contralateral to the AAV injection side (right) subtracted from turns ipsilateral to AAV injection side (left)] compared to saline administration. The Saline blue columns are not visible, because of a lack of rotation asymmetry. **D.** CNO (2 mg/kg) induced contralateral rotation asymmetry compared to saline administration. **A** and **B**: The box and whiskers plots show range and quartiles. The boxes extend from the 25th percentile to the 75th percentile, with a line at the median. The whiskers show the highest and the lowest values. Data (n=8) were analyzed using paired t-test. (**A**) 0-60 min and 120-180 min: *t*_(7)_ = 3.272, *p* = 0.0136, and *t*_(7)_ = 1.875, NS, respectively. (**B**) 0-60 min and 120-180 min: *t*_(7)_ = 3.173, *p* = 0.0157, and *t*_(7)_ = 4.465, *p* = 0.0029, respectively. \**p* ≤ 0.05, \*\**p* ≤ 0.01. **C** and **D**: Data (mean ± S.E.M, n=8) were analyzed using repeated-measures two-way ANOVA: (**C**) 0.1 mg/kg CLZ: effect of time, *F*_(11,154)_ = 2.34, *p* = 0.0110; effect of treatment, *F*_(1,154)_ = 2.59, *p* = 0.1297. ; interaction, *F*_(11,154)_ = 2.38, *p* = 0.0095. (**D**) 2 mg/kg CNO: effect of time, *F*_(11,154)_ = 2.16, *p* = 0.0194; effect of treatment, *F*_(1,154)_ = 3.26, *p* = 0.0926; interaction, *F*_(11,154)_ = 1.89, *p* = 0.044. In **C** and **D** *post hoc* comparison drug vs. saline: Bonferroni’s test, \**p* ≤ 0.05, \*\**p* ≤ 0.01.

### DREADD stimulation of left D1-SPNs alters fALFF bilaterally in multiple forebrain regions

The raw fMRI images show that there was no susceptibility artifact in AAV-injected site (**Supplementary Fig. 2**). In the CLZ-injected group (DREADD-CLZ), fALFF was bilaterally increased in cortical areas as compared to the control groups (**Fig. 4A**). These bilateral cortical areas included the somatosensory cortex (SC) and motor cortex (MC). fALFF was also increased in the left medial thalamic nuclei (ThN) and both left and right nucleus accumbens (NAc, **Fig. 4A**). A significant increase of fALFF was observed in left DS (DREADDs-expressing area) although it was a small spot **(Fig. 4A)**.

**Figure 4:**
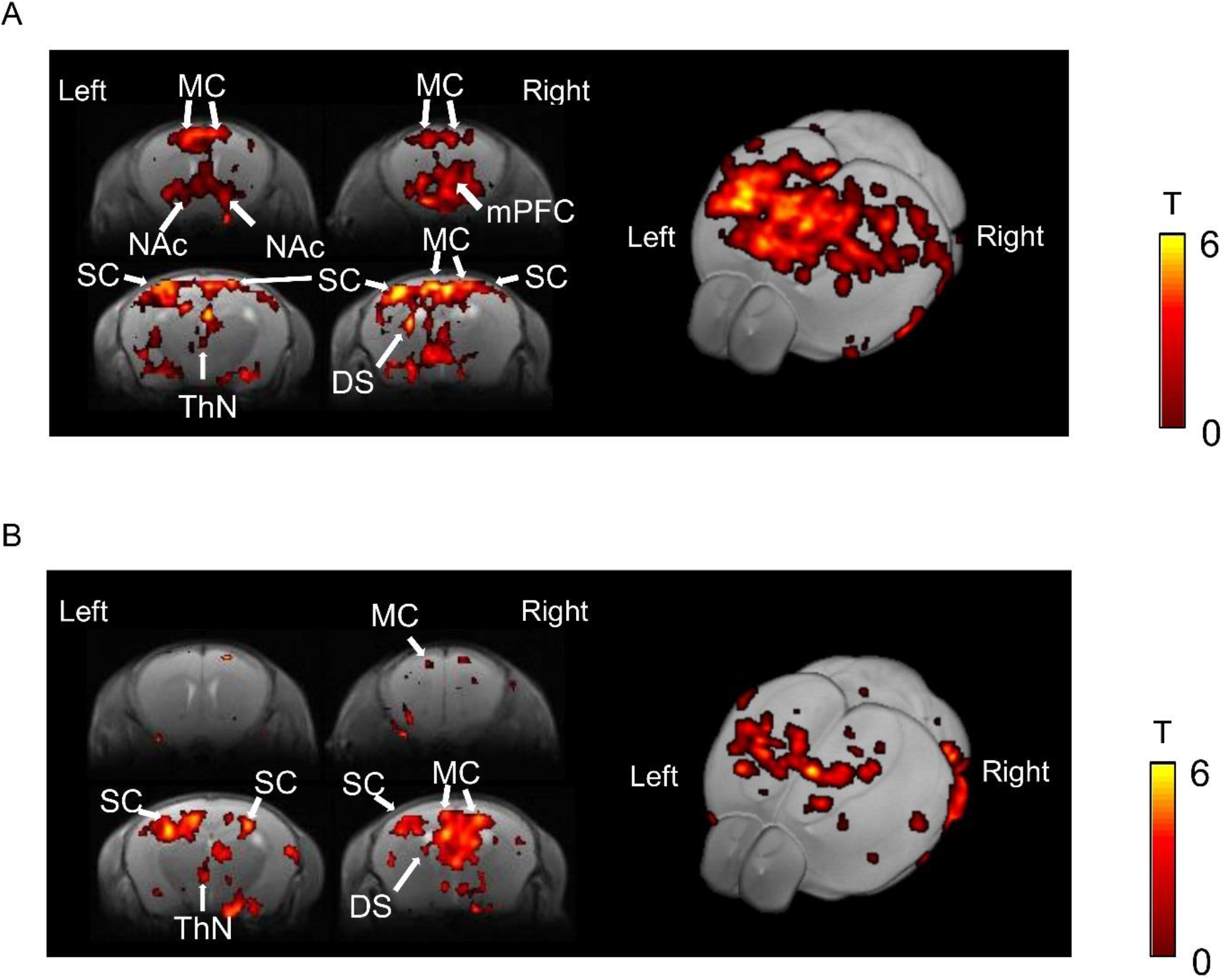
DREADD activation of left DS D1-SPNs increases fALFF bilaterally in forebrain. **A.** Significant increase in fALFF following CLZ injection compared to the control groups (p < 0.05, FDR at cluster level). Four coronal sections (+1.0, 0.5, −1.0 and −2.0 mm from Bregma respectively) are shown (left panels) as well as a 3-D reconstruction. Color bar, t-values. **B.** Significant increase in fALFF following CNO injection compared to the control groups (p < 0.05, FDR at cluster level). Color bar, t-values. mPFC, medial prefrontal cortex; SC, somatosensory cortex; MC, motor cortex; DS, dorsal striatum; NAc, nucleus accumbens; ThN, thalamic nuclei.

In the CNO-injected group (DREADD-CNO), fALFF was increased compared to the control group in the same regions (left DS, SC, MC, and left ThN) as in the DREADD-CLZ group, but with a smaller number of voxels than in the DREADD-CLZ group (**Fig. 4B**). In addition, no fALFF increase was observed in the NAc (**Fig. 4B**). Interestingly, the increase in fALFF was bilateral in many regions for both drugs and areas of increased fALFF were clearly larger in the DREADD-CLZ group than in the DREADD-CNO group.

### Alteration of the BLP in the contra-DS and bilateral motor cortex

To verify the results of fALFF, we investigated the BLP changes following saline or CLZ injection in unilateral DREADD-expressed mice. Notably, the total LFP in ipsilateral DS started to increase 10 min following CLZ injection, the increase becoming significant at 30 min (**Fig. 5A**). The delta and gamma BLP significantly increase at 45–51 min following the CLZ injection compared to saline injection (**Fig. 5B**). The gamma BLP increased in both left and right motor cortices at 45-51 min following CLZ injection compared to saline injection (**Fig. 5C** and **5D**). Saline injection did not alter the BLP in motor cortex at all frequency bands compared with before injection

**Figure 5:**
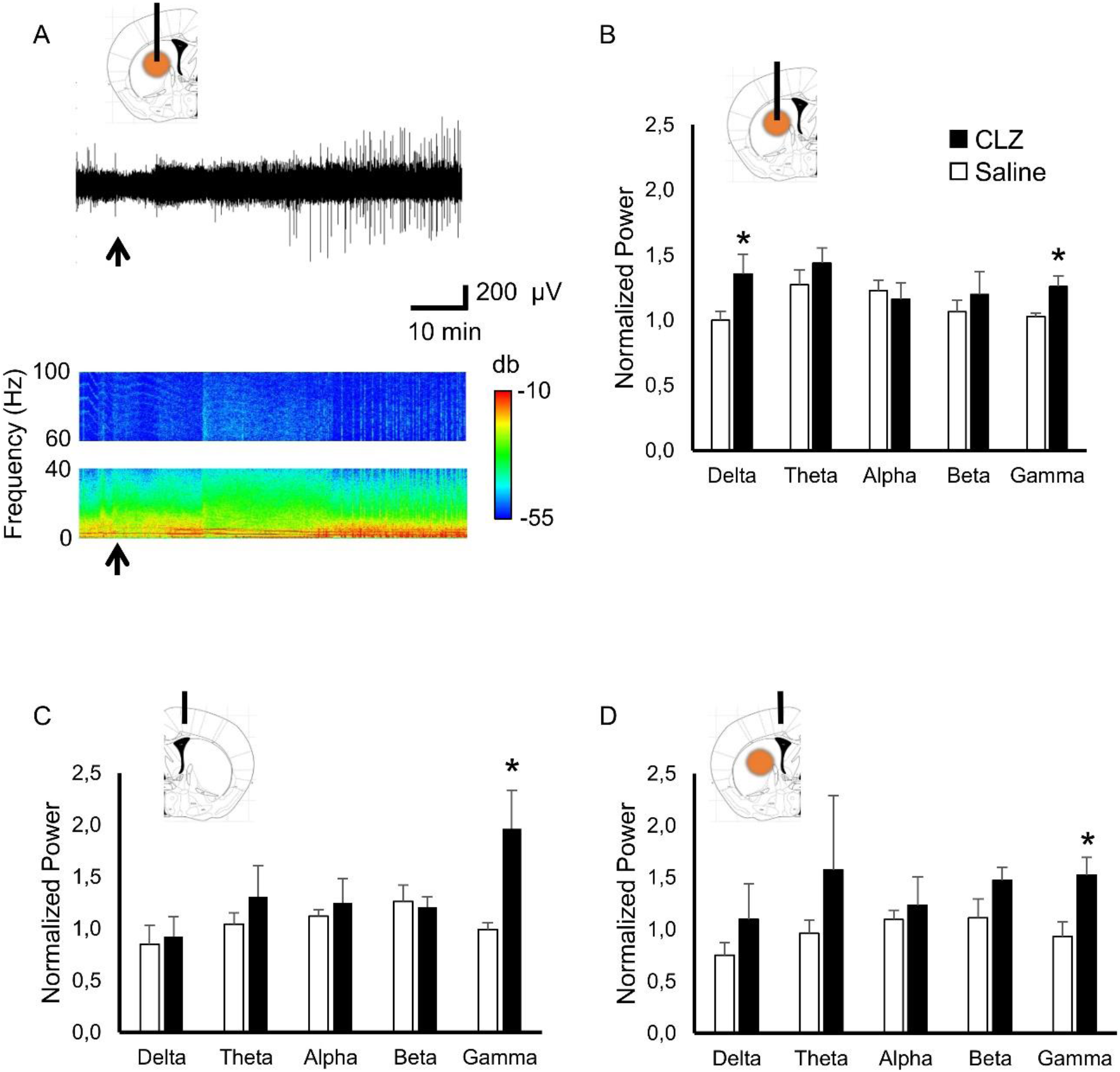
Effects of DREADD activation of left DS D1-SPNs on each BLP. **A.** (upper panel) LFP signals and (lower panel) power spectrum in the ipsilateral DS following the CLZ injection in a representative animal. Black arrows indicate the CLZ injection. Horizontal bar, time; vertical bar, LFP voltage; Color bar, power (db). **B.** Normalized power in each BLP (delta, theta, alpha, beta and gamma frequency bands) in the left DS. Each normalized power was averaged between 45 min and 51min. Data are expressed as mean ± SEM. *p < 0.05 compared with saline by paired t-test in each frequency bands. **C, D.** Normalized power in each BLP (delta, theta, alpha, beta and gamma frequency bands) in the right (C) and left (D) motor cortex. Each normalized power was averaged between 45 min and 51min. Data are expressed as mean ± SEM. *p < 0.05 compared with saline by paired t-test in each frequency bands.

## Discussion

In this study, using chemogenetic-fMRI, we observed that the prolonged unilateral activation of D1-SPNs in the DS affects spontaneous neuronal activity bilaterally at the time of behavioral effects. We also demonstrate that CLZ 0.1 mg/kg evokes behavioral effects more rapidly and alters fALFF in larger areas than CNO 2 mg/kg. Optogenetic fMRI (Lee et al., 2016) and chemogenetic fMRI have been used for the purpose of investigating the role of specific cell types in whole brain networks (Giorgi et al., 2017, Roelofs et al., 2017). Optogenetics can reversibly manipulate cell activity in the millisecond-order (Boyden et al., 2005), but this approach generates susceptibility artifacts due to the presence of the optical fibers and to the light that induces a vascular response in fMRI (Rungta RL et al., 2017, Christie et al., 2013, Schmid et al., 2017). DREADDs, in contrast, are ideally suited for prolonged modulation of cell activity in the range of minutes-hours (Armbruster et al., 2007, Alexander et al., 2009). Since chemogenetic-fMRI does not use optical fibers and light stimulation to activate specific cells, there is no susceptibility artifacts in the T2*-weighted images (see **Supplementary Fig. 2**). However, we should carefully consider the effective time window and possible unexpected DREADD-independent pharmacological effects of DREADD ligands.

### Different time-course of CNO and CLZ-induced behavioral changes

Previous reports showed a different time-course of CNO-induced cellular activation in the brain and peripheral organs. The time-course of CNO-evoked physiological effects in transgenic mice expressing hM3D (Gq) in pancreatic β cells correlates with the plasma concentration of CNO, with effects diminishing within 2 h (Guettier et al., 2009). In contrast, the behavioral and neuronal activity changes occur 30-40 min after CNO injection (0.3 mg/kg, i.p.) and these effects persist for 9 h (Alexander et al., 2009). In our study, we found that CNO started to change the rotational behavior from 30-40 min after CNO administration (2 mg/kg, i.p.) in the Gq-coupled hM3D DREADD-expressing mice and this alteration was still visible more than 2 h after injection. In contrast, CLZ induced a biased rotation that started 10 min after the injection, reached a maximum in 20-30 min and was not detectable 2 h after the injection. In addition, the rsfMRI at the time point of maximal behavioral changes showed that CLZ-evoked fALFF was increased in broader brain areas than after CNO, even though a 20-fold higher dose of CNO was used. A previous study showed that the activation of DREADD-expressing neurons induced by systemic CNO administration was mainly due to back-metabolized CLZ (Gomez et al., 2017). CNO (1 mg/kg, i.p.) is continuously metabolized to CLZ until 360 min (MacLaren et al., 2016), consistent with our results that behavioral changes continued more than 120 min after CNO injection. CLZ and N-desmethylclozapine, which is another pharmacologically active metabolite of CNO in the body, could possibly influence the behavior as indicated in previous studies (Schaber et al., 1998, MacLaren et al., 2016). Although low-doses of CLZ (0.05-0.1 mg/kg, i.p.) are reported to affect behavior in some conditioning (Ilg et al., 2018), we confirmed that CLZ (0.1 mg/kg, i.p.) did not affect the spontaneous locomotion in C57BL/6 mice (**Supplementary Fig. 3**). These results indicate that a low dose of CLZ instead of high doses of CNO was a more optimal treatment in terms of early pharmacological effect and short effective window.

### fALFF increase and neuronal activity

fALFF is related to the power of the low-frequency electrical current oscillations caused by excitatory neuronal activity (Ma et al., 2016, Zou et al., 2008). We observed increased fALFF in the basal ganglia-thalamo-cortical network including the medial part of the thalamus and several cortical areas including the motor/somatosensory cortices after CNO or CLZ administration, in accordance with the anatomical connectivity. Medial parts of the thalamus, including the ventromedial nuclei (VM), mediodorsal nuclei (MD) and the center median/parafascicular (CM/Pf) complex, have direct and indirect anatomical connection to the striatum and cortex (Groenewegen et al., 1994). The VM receives afferents from the basal ganglia (Herkenham, 1979, Kuramoto et al., 2011) and relays basal ganglia information to the cerebral cortex including motor areas (Kuramoto et al., 2015). The MD also receives projections from the basal ganglia, especially the internal segment of the globus pallidus and the ventral pallidum, to which striatal neurons project (Mitchell et al., 2013). The CM/Pf complex is connected to the entire striatal complex and the CM/Pf-striatal system is a functionally organized network that may broadly affect motor basal ganglia functions. The CM/Pf complex seems to play a pivotal role in PD because this area is severely degenerated in PD patients while deep brain stimulation of this nuclear group alleviates symptoms of PD (Smith et al., 2014).

Remarkably, the unilateral activation of the D1-SPNs in the DS increased fALFF bilaterally in the cortex and NAc. Other results demonstrating bilateral consequences of unilateral manipulations of basal ganglia circuits have also been reported in previous studies. The unilateral lesion of the thalamic VM nucleus induced a decrease of glucose metabolism in several structures of both cerebral hemispheres in basal conditions (Girault et al., 1985). In addition, bilateral effects depending on VM integrity were also observed following stimulation of the nigrothalamic pathway (Savaki et al., 1984), which mimics the activation of indirect pathway SPNs, although with less specificity. Dopaminergic projection from the ventral tegmental area to the NAc is thought to be almost exclusively unilateral (Nauta et al., 1978), although recent studies revealed some dopamine neurons project contralateral to their origin and showed cross-hemispheric synchronicity in dopamine signaling (Steinberg et al., 2014, Fox et al., 2016). Unilateral manipulations (lesioning, stimulation or pharmacological administration) of one nigrostriatal system reported to affect contralateral gene expressions, DA synthesis and turnover, and alterations in dopaminergic D1:D2 receptor interactions (Leviel et al., 1979, Chéramy et al., 1983, Schwarting et al., 1996, Roedter et al., 2001, Lieu et al., 2012, Yano et al., 2006, Hyde et al., 1994). Bilateral functional compensations were also observed by CBF-related tissue radioactivity after unilateral nigrostriatal damage (Yang et al., 2007). In addition, interhemispheric functional relationships of cortical neurons are also reported using optogenetics and voltage sensitive dye (Lim et al., 2012). fMRI studies of unilateral striatal neurons optogenetic manipulations also showed bilateral effects (Lee et al., 2016). Together with these reports, our results indicate that unilateral increased spontaneous activity of D1-SPNs in DS evokes a bilateral increase in neuronal activity in the prefrontal, somatosensory, and motor cortices. There appears to be functional and compensatory interdependence between right and left DS-networks.

In human, neuronal control of movement is largely lateralized, but several studies reported that bilateral striatal networks are involved in unilateral movement and may also play a role in the neural adaptations in PD. Unilateral GPi lesions can lead to a bilateral decrease in levodopa-induced dyskinesia (Baron et al., 1996, Lang et al., 1997), whereas unilateral high-frequency stimulation of the subthalamic nucleus (STN) can induce bilateral dyskinesia (Houeto et al., 2003, Brun et al., 2012). Unilateral STN stimulation have been reported to increase contralateral STN neuronal activity in PD patients (Novak et al., 2009, Walker et al., 2011).

Further study of rodent functional networks using DREADD technique and the availability of mouse models of brain disorders may offer new opportunities to study the large-scale functional implications of disease on these specific networks.

### fALFF and BLP alteration

In the present study, we confirm that the total LFP was increased in left DS (DREADD-expressing area) after CLZ injection. Furthermore, the gamma bands in the left DS and in both left and right motor cortex were significantly increased following the CLZ injection. Previous rodent and primate studies indicate that gamma band relates to the BOLD signals (Logothetis et al., 2001. Kayser et al., 2004. Niessing et al., 2005. Nir et al., 2007). In addition to gamma band, we observed the increase of delta power in the ipsilateral DS, but not in the ipsi and contralateral motor cortex. The glutamate levels were reported to be related to the gamma bands (Lally et al., 2014), while the effect of the dopaminergic and cholinergic neurons has been observed on low frequency fluctuation of LFP (Neske, 2016). The delta power of LFP in the ipsilateral DS was reduced by the disruption of dopamine in the resting state (Zhang et al., 2018). Although the mechanism of delta power increase in the ipsilateral DS is not clear, the change of dopaminergic or cholinergic neurons activation following the activation of striatal neurons may induce this delta power increase. In total, our results indicate that increased fALFF in the DS and motor cortex are derived from actual increase of neuronal activity.

## Conclusions

In conclusion, we successfully developed chemogenetic-fMRI system in the DS-network study and found bilateral effects following unilateral activation of DS. This method could help the comprehensive understanding of the whole brain network. However, the type and dosage of stimulant compounds for DREADDs should be carefully chosen considering the possibilities of DREADD-independent effects. This technique will contribute to the better interpretation of cell type-specific activity changes on MRI images, helping to evaluate the progression of neurological diseases and to unravel the effect of pharmacological agents upon acute and chronic treatment.

## Abbreviations

AAV: adeno-associated virus
ANOVA: analysis of variance
BLP: band limited power
CLZ: clozapine
CM: centromedial thalamic nucleus
CM/Pf: center median/parafascicular
CNO: clozapine-N-oxide
D1: dopamine D1 receptor (Drd1)
D1-SPN: Drd1-expressing SPN
DREADD: designer receptor exclusively activated by designer drugs
GPi: internal globus pallidus
D1-Cre: Cre under the control of *Drd1* promoter
DS: dorsal striatum
fALFF: fractional amplitude of low frequency fluctuation
GPe: external globus pallidus
hM3D: human M3 muscarinic DREADD
LFP: local field potential
MC: motor cortex
MD: mediodorsal thalamic nucleus
MRI: magnetic resonance imaging
NAc: nucleus accumbens
PD: Parkinson’s disease
RARE: rapid acquisition with relaxation enhancement
ROI: region of interest
rsfMRI: resting state functional magnetic resonance imaging
SC: somatosensory cortex
SPN: spiny projection neuron
STN: subthalamic nucleus
ThN: thalamic nuclei
VM: ventromedial nuclei

## Acknowledgments

Research in the laboratory of JAG was supported by Inserm and Sorbonne University and by grants from ERC AIG-250349, *Fondation pour la Recherche Médicale* (FRM) and ANR (Epitraces). The project is also supported by *Fondation de France* (grant number: 00086313, to DH). Equipment at the IFM was funded in part by *Fondation pour la recherche sur le cerveau* (FRC) and Rotary *Espoir en Tête*, and by DIM *Cerveau et Pensée Région Ile-de-France*. YN was recipient of a Uehara Memorial Foundation fellowship and a Fyssen Foundation fellowship. Equipment of the electrophysiology was funded by L’Idex Paris-Saclay.

---

**Supplementary Figure1.**
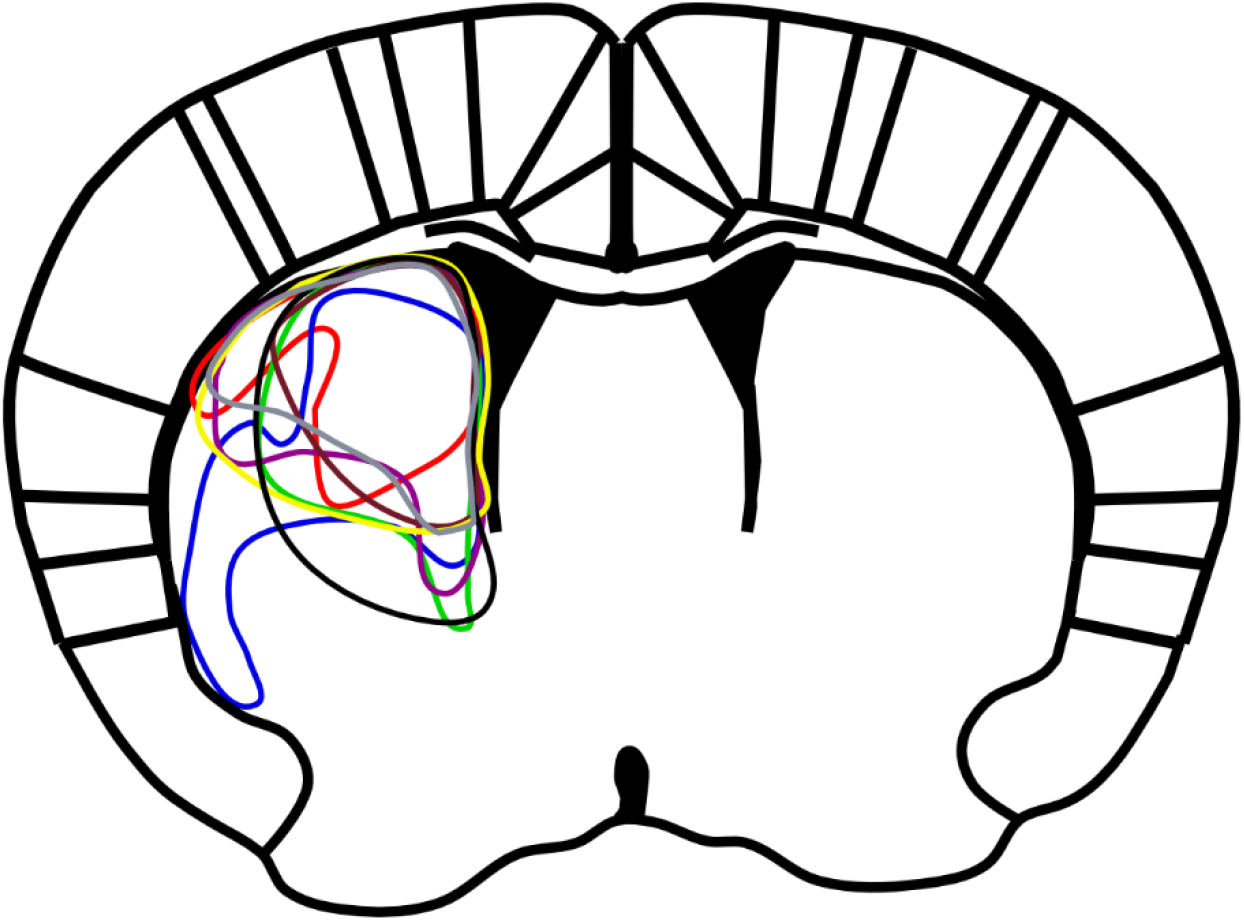
DREADD expression area of each mouse. Tracing the extention of mCherry expression in dorsal striatum of unilaterally injected D1-Cre mice (n = 8)

**Supplementary Figure2:**
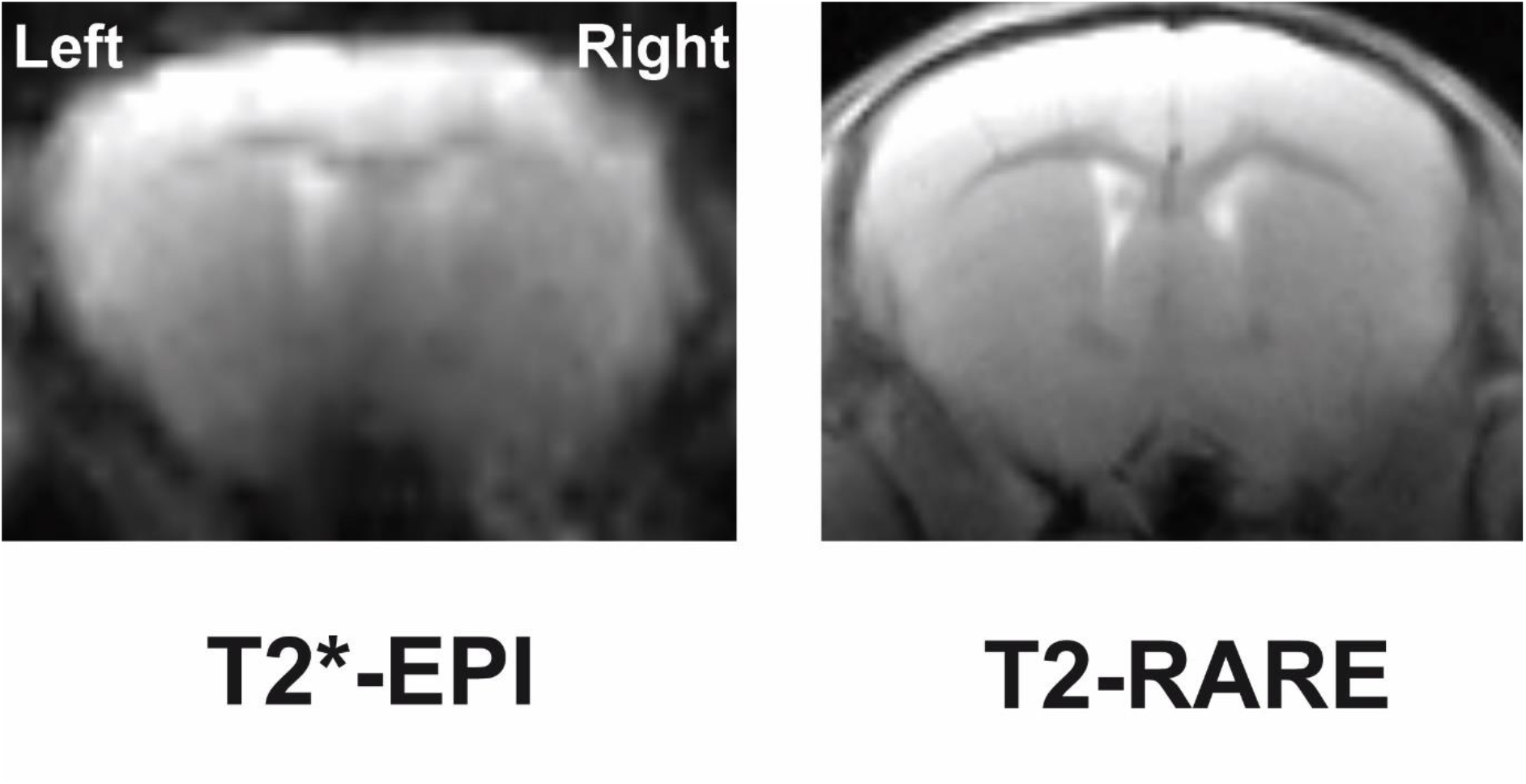
Raw images of T2*-EPI and T2-RARE in the same mouse (+0.3 mm from Bregma).

**Supplementary Figure3.**
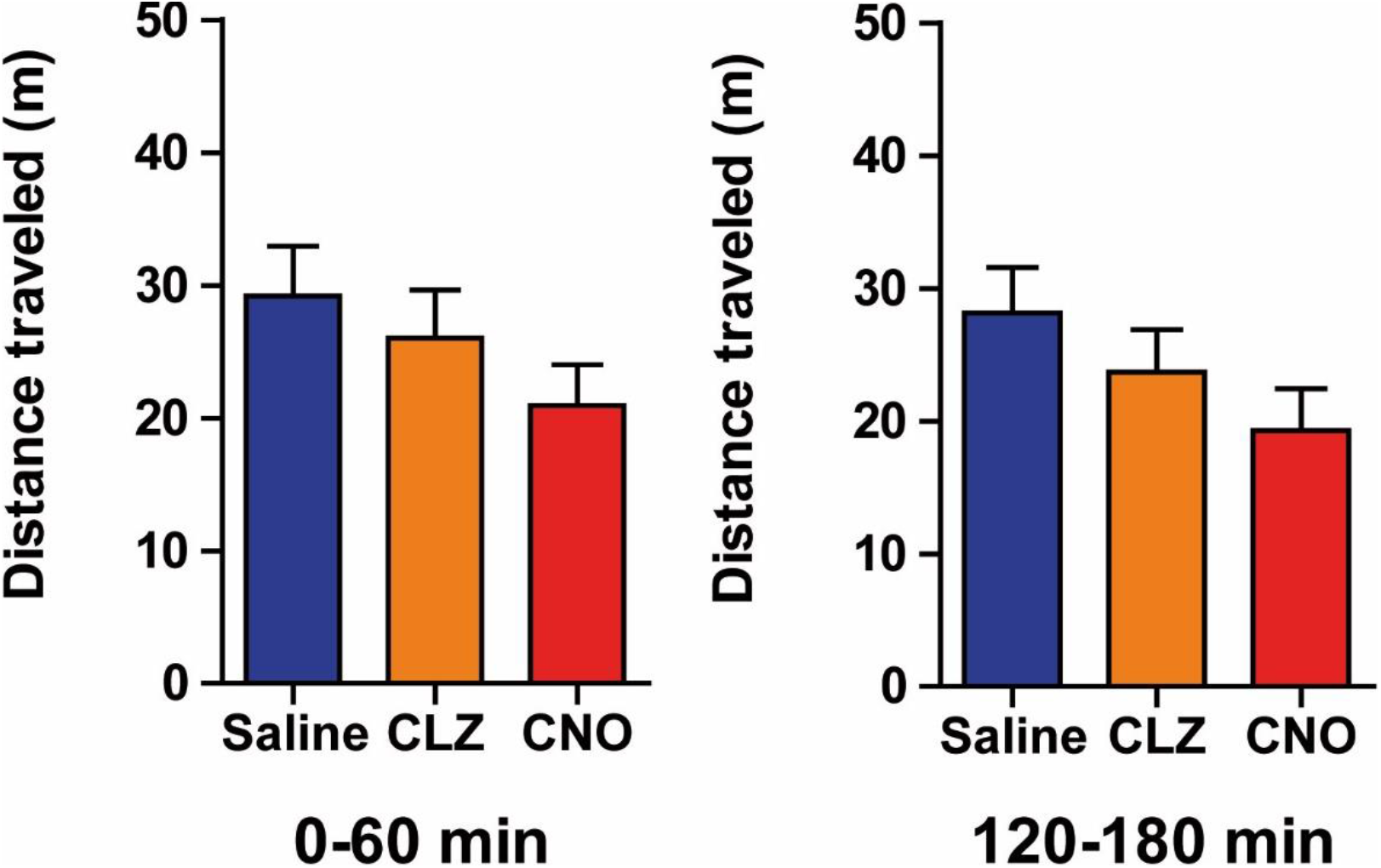
CNO and CLZ did not affect the basal locomotion of C57BL/6J mice. Mice were placed in a cylinder, injected with CNO, clozapine (CLZ) or saline and their locomotion were measured during the periods indicated in Fig. 1D. **A.** CLZ (0.1 mg/kg) and CNO (2 mg/kg) did not change locomotor activity 0 – 60 min.(**A**) and 120 - 180 min (**B**) after injection compared to saline administration. Data (mean ± S.E.M, n=8) were analyzed using one-way ANOVA: (**A**) *F*_(2,21)_ = 1.411, *p* = 0.2661, NS. (**B**) *F*_(2,21)_ = 1.836, *p* = 0.1841, NS.

